# Pan-cancer characterisation of microRNA with hallmarks of cancer reveals role of microRNA-mediated downregulation of tumour suppressor genes

**DOI:** 10.1101/238675

**Authors:** Andrew Dhawan, Jacob G. Scott, Adrian L. Harris, Francesca M. Buffa

**Affiliations:** Department of Oncology, University of Oxford, Oxford, United Kingdom; Translational Hematology and Radiology, Cleveland Clinic, Cleveland, United States

## Abstract

microRNA are key regulators of the human transcriptome across a number of diverse biological processes, such as development, aging, and cancer, where particular miRNA have been identified as tumour suppressive and oncogenic. In this work, we sought to elucidate, in a comprehensive manner, across 15 epithelial cancer types comprising 7,316 clinical samples from the Cancer Genome Atlas, the association of miRNA expression and target regulation with the phenotypic hallmarks of cancer. Utilising penalized regression techniques to integrate transcriptomic, methylation and mutation data, we find evidence for a complex map of interactions underlying the relationship of miRNA regulation and the hallmarks of cancer. This highlighted high redundancy for the oncomiR-1 cluster of oncogenic miRNAs, in particular hsa-miR-17-5p. In addition, we reveal extensive miRNA regulation of tumour suppressor genes such as PTEN, FAT4, and CDK12, uncovering an alternative mechanism of repression in the absence of mutation, methylation or copy number changes.

---

The hallmarks of cancer very clearly outline the major phenotypic changes underlying the oncogenic process [24, 25]. These changes characterise cancer as a disease, and may define actionable targets for therapeutic intervention. Since the definition of these characteristic hallmarks in 2001 [24], and the subsequent ‘genomic revolution’ that has occurred in the field of cancer biology, multiple groups have proposed gene expression signatures as biomarkers of these phenotypic hallmarks [26, 47, 53]. These gene signatures generally consist of a set of tens to several hundred coding genes, for which a summary metric of their collective expression is associated with a known hallmark, and may help with defining therapeutic strategies [3]. Encapsulated within this methodology and these signatures is a vast amount of biological discovery for particular genes implicated in the development and progression of these hallmarks. However, since the more recent publication of the updated hallmarks in 2011 [25], there has been a second revolution in the field of genomics; namely, the discovery of the diverse, critical roles of non-coding RNAs in cancer.

Previously thought to be ‘junk DNA,’ non-coding RNA are those RNA derived from DNA that do not code for proteins, and consists of a diverse family of evolutionarily conserved species, including long non-coding RNA (lncRNA), circular RNA(circRNA), and microRNA(miRNA), among others [23, 40, 41]. Much effort has focused on the characterisation of these non-coding RNA, and early work has shown that these species, particularly miRNA, are involved in a number of cellular developmental, and differentiation processes [50]. In addition, miRNA have been implicated in a number of human diseases, ranging from diabetes to cancer, and in oncology, recent work has led to the discovery of tumour-suppressive and oncogenic miRNA [7,42,44,49]. miRNA exert their function within the cell primarily as repressors of protein production, functioning as post-transcriptional regulators of mRNA, inhibiting translation or encouraging transcript degradation. miRNA exert their effects by complementary base-pair binding to a short 7-8 nucleotide ‘seed region’ typically located on the 3’ untranslated region of the messenger RNA which they inhibit [40]. A single miRNA is thought to able to exert its repressive effects on hundreds to thousands of transcripts, meaning that specific miRNA may have very wide-ranging and fast-acting effects on cellular phenotype [40]. Despite this potential, due to the highly variable effect on the single target transcripts and the many factors involved in post-transcriptional gene regulation in addition to miRNA, the repressive signal on their targets, both validated targets and predicted targets by sequence complementarity, remains challenging to detect in clinical datasets [6]. As a result, behavioural characterisation of miRNA has been progressing at a slow rate, with studies focusing on changes induced by a single miRNA or small families of miRNA, without any efforts for large-scale characterisation.

A further complicating factor with respect to the study of miRNAs is the relative promiscuity of their targets [36]. A given miRNA may have thousands of targets, with many experimentally verified, but often these targets possess significant differences in function [54]. This has led to an almost paradoxical finding about the effects of miRNAs, in that a single miRNA may theoretically exert effects in opposing directions within the cell [54]. This paradox is resolved by the observation that miRNA likely play different roles depending on the environment in which they are expressed [10,20,36]. Therefore, in addition to the challenge of measuring the repressive effect of miRNAs within a transcriptome, the effect of a miRNA on a transcriptome may vary massively, depending on the relative abundance of each of its targets. That is, a miRNA may only repress targets to which it is able to bind, and this requires the presence of the target in a detectable concentration compared to all others [14]. This means that the effect of a miRNA on phenotype can only be observed in samples for which the transcriptomes are comparable in the expression of the key targets in consideration, and such effects are highly context-dependent.

In this work, we show how this context-dependent action can be exploited to gain high confidence predictions uncovering known and unknown associations with miRNA and phenotype. Through the classification of tumour transcriptomes by gene expression signatures, we uncover the diverse roles of miRNAs in regulating the hallmarks of cancer.

Our results point towards a scenario wherein the trancriptome of the cancer cell, known to be driven by dysregulation of tumour suppressor genes and oncogenes, is heavily regulated by miRNAs. We show that predicted miRNA-target associations that retain significance across multiple cancer types involve a number of critical tumour suppressor genes and oncogenes. Study of these tumour suppressor genes yields novel conclusions about their regulation, particularly with respect to their repression by miRNA, methylation and mutation, and the exclusivity of the occurrence of these modes of regulation across human cancers.

## Results

### Evaluation of Hallmark gene signatures across cancers

The first prerequisite to our study was to identify suitable biomarkers to infer cancer phenotype. In order to achieve this, we chose 24 previously identified gene expression signatures (Supplementary S1) that have already been shown to be representative for a wide number of samples, and a number of fundamental phenotypic properties, with the hopes of alleviating issues related to highly tissue-specific expression patterns. With this in mind, we applied *sigQC*, an R package encapsulating a robust methodology for the evaluation of gene signatures on various datasets for the basic statistical properties underlying their applicability [16]. We ran this package on all combinations of 15 datasets and 24 signatures considered in this study, and tested the consistency of signature performance across cancer types, giving confidence in the application of the signatures to these datasets. All summary plots from the *sigQC* quality control protocol are presented in Supplementary Section S2. Each of the signatures considered over the 15 epithelial cancer datasets showed good applicability, strong signature gene expression, moderate-strong compactness, and good gene signature score variability, as well as strong autocorrelation of signature metrics. The previous validation of these signatures, and our study-specific quality control results, justify our subsequent use of these signatures in a pan-cancer manner, to identify conserved associations of miRNA and signature gene expression across tissue types.

### Hallmark gene signatures association analysis reveals a complex pan-cancer miRNA regulatory network

To determine the association of gene signatures to miRNA expression, we set the signature score (see Online Methods) for each signature equal to a linear model consisting of all miRNAs showing at least moderate univariate predictive ability for the signature summary score, as depicted in Figure 1a. Multivariable linear modelling with L1/L2 penalized regression optimized by cross-validation was used as previously described [6] to identify the miRNAs which showed the greatest predictive ability for each hallmark signature score across the cancer types considered, thereby identifying those miRNA common to the gene signature across tumour types (see Online Methods). An example of the values for miRNA coefficients across cancer types following the model fitting is depicted in Figure 1b. miRNAs were then ranked based on their final model coefficient (reflective of the strength of association to the signature), and miRNAs consistently ranking highly as positive predictors of a given hallmark signature across cancer types were aggregated, from which statistically significant miRNAs were isolated with the rank product test (signature-associated miRNAs). Likewise, for each gene signature, the miRNAs most consistently ranked as strong negative predictors of signature score across cancer types were aggregated by a rank-based methodology (negatively signature-associated miRNA), as depicted in Figure 1c. This analysis reveals both many known and unknown significant associations between miRNA and gene signature scores, facilitating an understanding of the miRNA involved with hallmark phenotypes, providing both novel hypotheses, and adding to evidence for existing ones.

**Figure 1.**
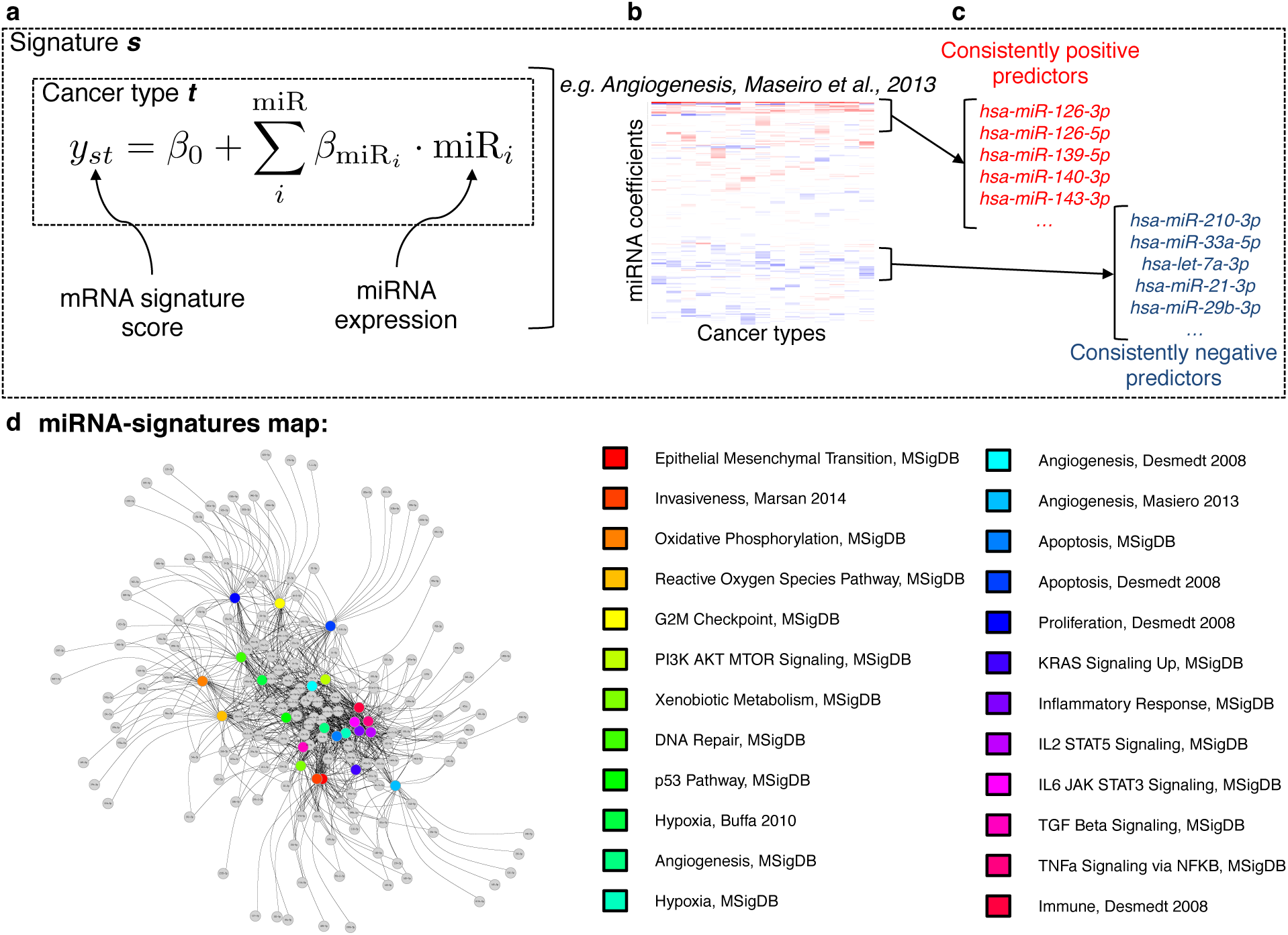
Overview of approach used to identify hallmarks-associated miRNA. (**a**) Overview of the linear model used in the fitting, for each gene signature and cancer type under consideration, (**b**) Example of a heatmap depicting the values of the coefficients identified for the miRNA predictors (rows), across cancer types (columns) for our previously developed angiogenesis signature [39]. (**c**) Consistently positive and negatively ranking miRNA coefficients, identified as statistically significant by the rank product statistic, are taken as the positive and negative hallmark-associated miRNA for each hallmark signature. (**d**) Network ‘map’ of signatures (coloured nodes) and their positively associated miRNA (grey nodes), connected by edges when an association was found, highlighting strong interconnectivity between distinct molecular signatures.

To verify the validity of these predictions, we considered the example case of miRNAs found to associate significantly with the hypoxia signatures considered. Hypoxia is one of the most studied microenvironmental perturbations in the context of miRNA regulation, and one with a very well-defined pathway, controlled largely by a single transcription factor, HIF-1*α* [48]. Taking the intersection of the sets of miRNAs found to associate positively with the two previously validated hypoxia gene signatures (Hypoxia, Buffa et al. [5], and Hypoxia, MSigDb [34]), we obtained high confidence predictions for hypoxia-associated miRNAs.

As shown in the Tables associated with Supplementary S3, this analysis reveals that many of the miRNAs found to be commonly associated with both hypoxia gene signatures have been previuosly identified as hypoxia regulated. High confidence predictions are made for: hsa-miR-210-3p [8], −21-3p, −21-5p, −23a-5p, −23a-3p, −24-3p, −24-2-5p, −27a-5p, [31], let-7e-5p, let-7e-3p [11], −22-5p, −22-3p [57]. This analysis also suggests significant, pan-cancer, potential roles for other members of the let-7 family of miRNAs in hypoxia; namely, let-7b-5p, let-7b-3p, let-7d-5p, let-7d-3p, as well as hsa-miR-223-3p, −18a-5p, and −28-3p, which have potentially escaped the notice of other approaches.

In the context of all gene signatures considered, we identify a global underlying ‘map’ connecting each miRNA to each gene signature with which we have found an association. As shown in Figure 1d, this is a highly interconnected and complex network, with the conservation of a core set of miRNAs shared across the hallmarks of cancer. A similar analysis reveals an analogous result for the miRNA-hallmarks network for the miRNA negatively associated with both signatures, as described in Supplementary Section S4. To validate the reproducibility of these results, we rebuilt the signature-miRNA linear model using a large independent dataset, the Metabric breast cancer cohort [13]. The miRNA identified as positively and negatively associated with gene signatures in this dataset show highly significant concordance over a majority of signatures with the corresponding miRNAs identified from analysis of the TCGA dataset (Supplementary Figure **??**, Section S5).

### Multiple members of the same miRNA family display opposite tumour suppressor and oncogenic behaviour

Subsets of miRNAs that typically share common, evolutionarily-conserved sequences or functional motifs in their sequences are grouped into families [28, 29]. Interestingly, grouping the miRNAs found to be significantly upregulated and significantly downregulated in association with each of the gene signatures considered reveals that a number of miRNAs from the same families are present in different sets. That is, as summarised in Supplementary Section S6, many of the same miRNA families contain a significant number of miRNAs, some of which are positively and others negatively associated across gene signatures for the hallmarks of cancer. In particular, the miR-17/17-5p/20ab/20b-5p/93/106ab/427/518a-3p/519 and let-7/98/4458/4500 families have multiple members across signatures both in statistically significant positive and negative associations. This highlights once more the context-dependent nature of miRNA regulation, and the potentially antagonistic behaviours of miRNAs when grouped by family, supporting previous findings. Here, we argue that such a grouping does not necessarily reflect conserved function in the different tumour tissues, and we highlight that an additional context-dependent functional miRNA classification uncovering key functional associations is desirable.

### Hallmarks-associated miRNA targets are significantly enriched for tumour suppressor genes

Starting from a list of positively associated miRNA with each gene signature, we aimed to identify which predicted miRNA-target pairs showed strong evidence of negative regulation across cancer types. The union of five miRNA target prediction algorithms, as implemented by the Bioconductor package miR-NAtap was used [45], with a minimum number of two sources required to be included in the analysis (see Methods). We considered only the miRNA and predicted target mRNA pairs for which there was a statistically significant negative Spearman correlation of expression across at least 5 cancer types, and used a rank-product test to identify the miRNA-target pairs showing consistency across cancer types (Figure 2a). As depicted by the process in Figure 2b-c, analysis of these significant miRNA-target pairs revealed a strong enrichment for tumour suppressor genes (as defined by the COSMIC database list of 141 TSG), as might be expected for miRNA associated with oncogenic processes (p = 0.0006, two-sided Fisher’s exact test). To further test the significance of increased number of TSG repressed by the signature-associated miRNA, a bootstrap resampling-based approach (see Methods), was devised. From all expressed miRNA across cancer types that could have been chosen as signature-associated miRNA, random lists of the same length as the number of signature-associated miRNA were chosen, and, via an analogous approach as above, the number of repressed TSG for these miRNA was determined. Repeating this resampling 1000x, the probability that 21 or more TSG were repressed by the chosen miRNA was *p* = 0.017, again suggesting strong significance in the enrichment for TSG among the repressed targets of the signature-associated miRNA. This suggests that miRNA-mediated repression of tumour suppressor genes may be relatively common, significant, and associated with the phenotypic hallmarks of cancer.

**Figure 2.**
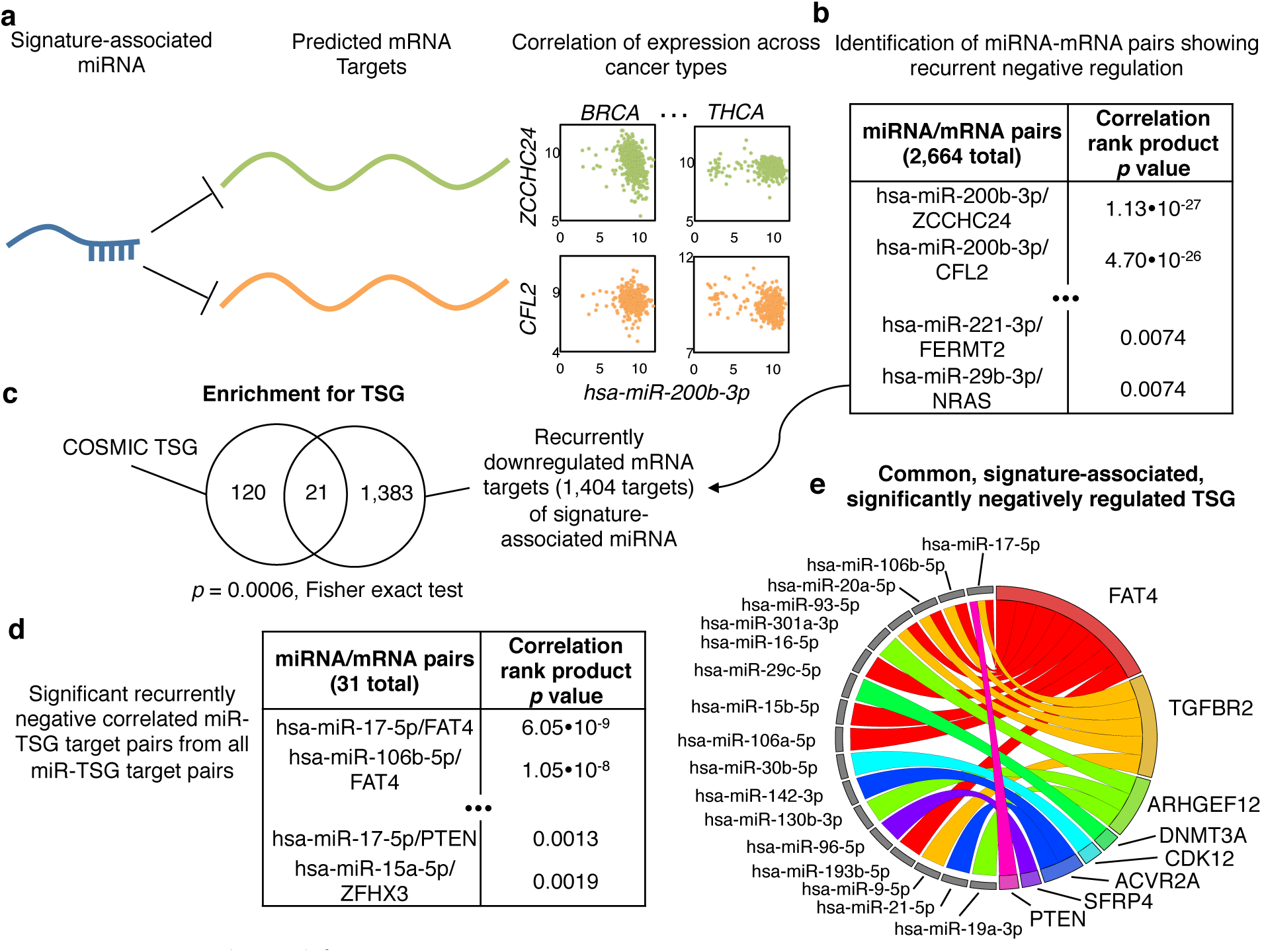
Approach used for interpreting miRNA-target interactions. (**a**) First, miRNA-target pairs for each positively associated hallmark-associated miRNA were identified, and the correlation between these was determined. (**b**) Next, the correlations across cancer types were aggregated, and those identified as consistently negative-ranking were identified with the rank product statistic. (**c**) Among this list of miRNA-mRNA target pairs, there was highly significant enrichment for tumour suppressor genes, as identified by the Fisher exact test. (**d**) The same procedure as described in (**a**) and (**b**) was repeated for all miRNA and all predicted target TSG pairs. (**e**) From the lists identified in (**b**) and (**d**), we identified those miRNA-TSG pairs in common, and plot their interactions on a circos plot, showing the repressive actions of each miRNA on its predicted target TSG.

A different picture emerged upon repeating this analysis for oncogenes, and for the miRNAs found to be significantly negatively associated with one or more hallmark signature. We identified 1283 significantly anti-correlated miRNA-target pairs for these downregulated hallmark-associated miRNAs. Likewise, analysing all predicted miRNA-oncogene interactions among the 231 COSMIC oncogenes, there were only 2 showing significant anticorrelation across tumour types with their predicted target miRNA (ESR1 and ABL2). Taking the intersection of these lists of 2 COSMIC oncogenes and the 1283 miRNA-oncogene pairs associated with gene signatures identified only ESR1 (interacting with miR-18a-5p and miR-130b-3p) in common (*p* = 1.2 · 10^−5^, Fisher’s exact test). This suggests that ESR1, estrogen receptor alpha, may play a significant role across the hallmarks of cancer, and de-repression by reduction of its miRNA-mediated repression may play a role in cancer phenotype, and ultimately, oncogenesis [35, 52]. On the other hand, this result is also a strong negative control for our analysis, and it concurs in supporting the common oncogenic role of miRNAs via co-ordinated repression of tumour suppressor genes.

### A core set of tumour suppressor genes are associated with the hallmark gene signatures across cancer types

Next, we asked whether our results could be biased by the initial selection of miRNA, namely the ones associated with the cancer hallmarks. To answer this, we conducted a complementary analysis, namely we sought to determine which of the miRNA-mediated tumour suppressor genes showed significance in downregulation, in the context of all other tumour suppressor genes. Thus, we repeated the previous analysis extended to all predicted miRNA-TSG pairs, considering again the significant associations across at least 5 cancer types, and then collated with a rank product test, as summarised by Figure 2d. Considering the miRNA-TSG pairs found to be of significance in both analyses from Figures 2c and d, we identified a set of 22 miRNA-TSG pairs, comprising 8 TSG (FAT4, TGFBR2, ARHGEF12, DNMT3A, CDK12, ACVR2A, SFRP4, and PTEN) and 17 miRNAs in Figure 2e, in common. We show also that the miRNA found to be associated to each of these TSG are, in many cancer types, expressed at significantly higher levels in wildtype cases for the associated TSG, across multiple tumour types (Supplementary Figure **??**, Section S7). These results show that for these tumour suppressor genes, i) miRNA-TSG interactions show significance across cancer types, and more so than all other TSG considered, ii) miRNA-TSG interactions show strong associations with the phenotypic hallmarks of cancer, and iii) miRNA-TSG interactions may show increased importance in cases with wild-type TSG. Importantly, the conserved miRNA-TSG regulation across cancer types reveals this as a potential new common epigenetic mechanism, alternative to genetic mutations, to achieve functional inhibition of TSGs in cancer.

### The action of hallmarks-associated miRNAs shows cancer context-dependency

The presented analysis highlights the action of miRNA in cancer. However, to further understand if this was cancer-specific, we sought to determine whether similar conclusions could be reached when analysing non-tumour tissues. Starting from the associated adjacent normal tissue datasets from TCGA for tissue types with at least 20 samples for both miRNA and mRNA expression (BRCA, UCEC, HNSC, KIRC, LUAD, and BLCA), we fitted a linear model for gene signature score as a function of all miRNA, for each signature, in each of the 6 tissue types. Aggregating coefficients across tissue types, we found that while a highly significant number of miRNA associated with the gene signature scores across tissue types are the same as uncovered for the cancer tissues, there are significant differences. Across signatures, an overlap of on average 54% was observed for signature-associated miRNA, showing high statistical significance for miRNA positively and negatively associated with signatures (*p* < 10^−19^ in all cases, by Fisher’s exact test).

Examining the targets of these positively signature-associated miRNA from normal tissues, we identified 233 recurrently negatively correlated miRNA-target pairs, of which two contain miRNA-TSG pairs (CEBPA and NCOA4). However, this overlap of the 142 unique genes among the 233 miRNA-target pairs with the 141 COSMIC tumour suppressor genes does not show significance, and may be due to chance alone (*p* = 0.26 by Fisher’s exact test). Thus, while the biology captured by the phenotypes of the gene signatures may be consistent, more than chance alone would predict, between tumour and normal samples, the resultant miRNA-target interactions are significantly different, and miRNA-TSG enrichment is not retained among normal tissue samples, highlighting the context dependency of these associations.

### Analysis of modes of regulation confirms that copy number and mutational status are key determinants of TSG expression

With a set of TSG purported to be significantly regulated by miRNA in relation to phenotype identified, we next sought to characterise the determinants of their expression. In particular, we consider an approach integrating multiple lines of genomic information; namely, methylation status, copy number, miRNA expression, and mutational status (see Methods), with the linear model depicted in Figure 3a. Notably, when considering the impact of miRNA in this model, we considered all reported miRNA to potentially discover novel miRNA-target interactions. We then fit this model with penalised linear regression over the various cancer types, and then subsequently aggregated coefficients by the rank product statistic to identify recurrently positive and negative predictors across cancer types, for each of the 8 tumour suppressor genes identified in Figure 2e. This analysis yields both expected results, such as the important positive predictive role of copy number for each of the tumour suppressor genes, as seen in the left panel of Figure 3b, and novel associations, such as the positive association of many miRNA, and some methylation probes with TSG expression in some cases. These miRNA may be co-expressed for a variety of reasons, such as competitive interactions, repression of repressors of the TSG, or a nearby genomic locus, though penalised regression minimises the effect of co-location because of the inclusion of copy number as a covariate.

**Figure 3.**
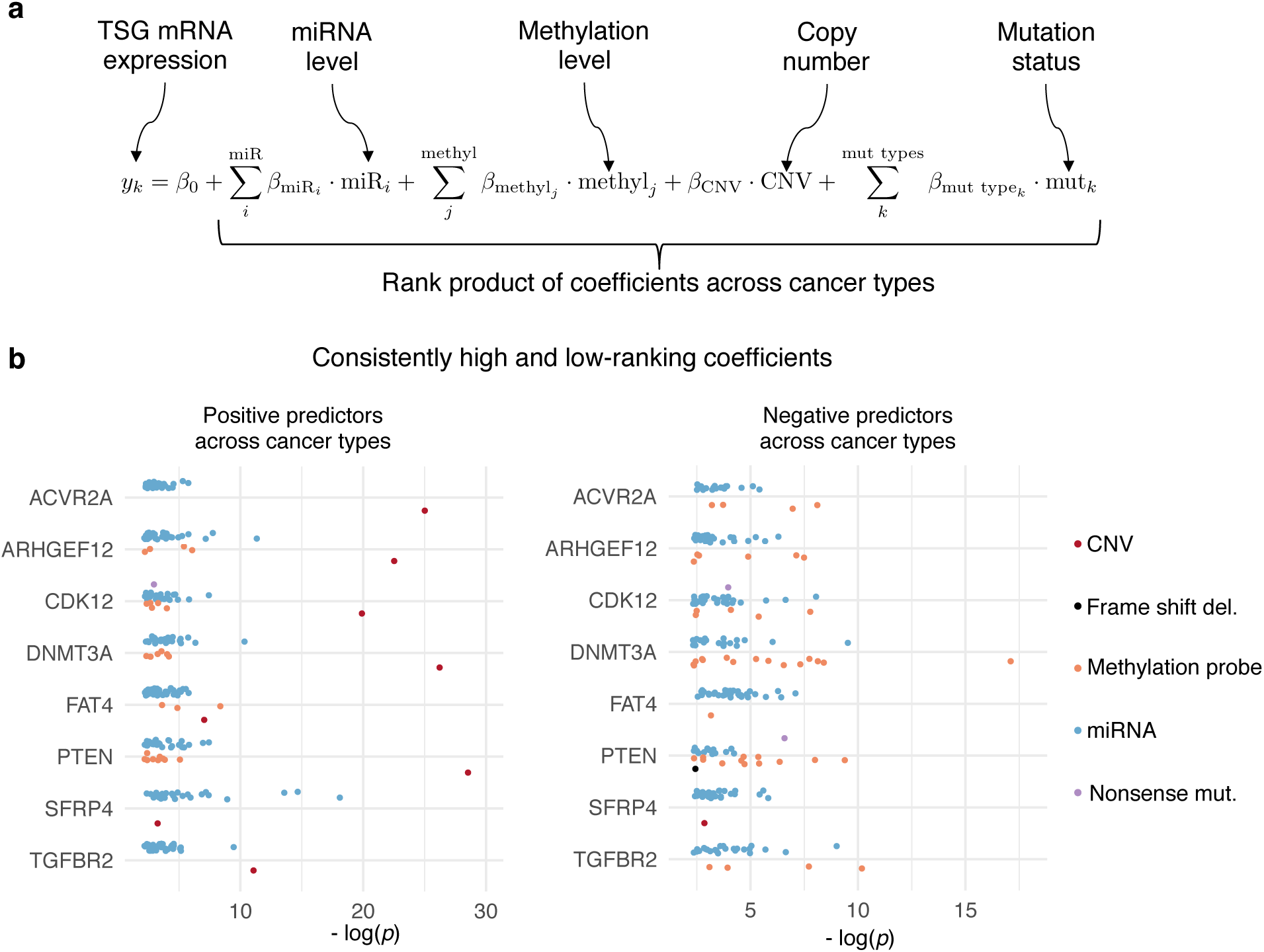
Approach used in determining the regulation of each TSG identified as potentially significantly miRNA-regulated. (**a**) The linear model used whilst determining predictors of TSG mRNA expression. (**b**) Model coefficients were aggregated across cancer types with the rank product statistic, and those identified as statistically significant positive and negative predictors are depicted alongside the - log of their rank product p-value.

Likewise, the identified modes of negative regulation give expected results, with non-sense mutations and frame shift deletions consistently negatively associated with TSG mRNA expression. Further, because this analysis was done with all miRNA, and not just those predicted to have a given TSG target, these results may implicate novel miRNA-TSG interactions. The complete rank product tables and all autocorrelation matrices can be found in Supplementary Section S8.

### PTEN, FAT4, and CDK12 tumour suppressor genes show exclusive regulation by either miRNA, promoter methylation or mutation across cancer types

Once the modes of regulation and their relative importance was established (Figure 3), we sought to determine the relative occurrence of each of these modes of regulation. We identified which negative regulators co-occurred with each other as synergistic repressors, and conversely which were exclusive repressors (Figure 4a). A cursory analysis of autocorrelation heatmaps (e.g. Figure 4a) revealed that in some cases, the regulation by miRNA appeared to be exclusive from the regulation by methylation probes. A full series of heatmaps for all cancer types considered and all tumour suppressor genes with their associated negative regulators identified is presented in Supplementary Section S9, Figures **??**- **??**, and for an independent dataset in Figure **??**, details described in Online Methods. These results suggest that TSG expression can be altered by either miRNA or methylation, in addition to deletion or mutation, in a ‘BRCA-ness’-like phenomenon [43]. To characterise this, we devised a bootstrap resampling based approach (see Online Methods), to determine significance of the difference in co-correlation between the miRNA and the methylation probes themselves, and then with each other. For each cancer type, we calculated the significance value of this proportion (Figure 4b), and from this analysis, it arose immediately that for each of the TSG considered, there are tumour types in which the regulation is consistently exclusive. Further, it also arose that across multiple cancer types, three key tumour suppressor genes, PTEN, FAT4, and CDK12, consistently tend towards exclusivity in their regulation, lending support for the importance of miRNA-based regulation of these genes. We further use the identified negatively associated miRNA and methylation probes, along with mutation status, to define subgroups of samples, for which we show decreased TSG expression in the subgroups with high expression of these miRNA or high methylation of these probes, in Figures **??**- **??** in Supplementary section S10. Further, we show that the miRNA-high and highly methylated samples have transcriptomes altered in a similar manner as in TSG mutated cases, via an analysis of differentially expressed genes in both cases, with significantly positively associated fold changes across cases, in Figures **??**- **??** in Supplementary Section S10.

**Figure 4.**
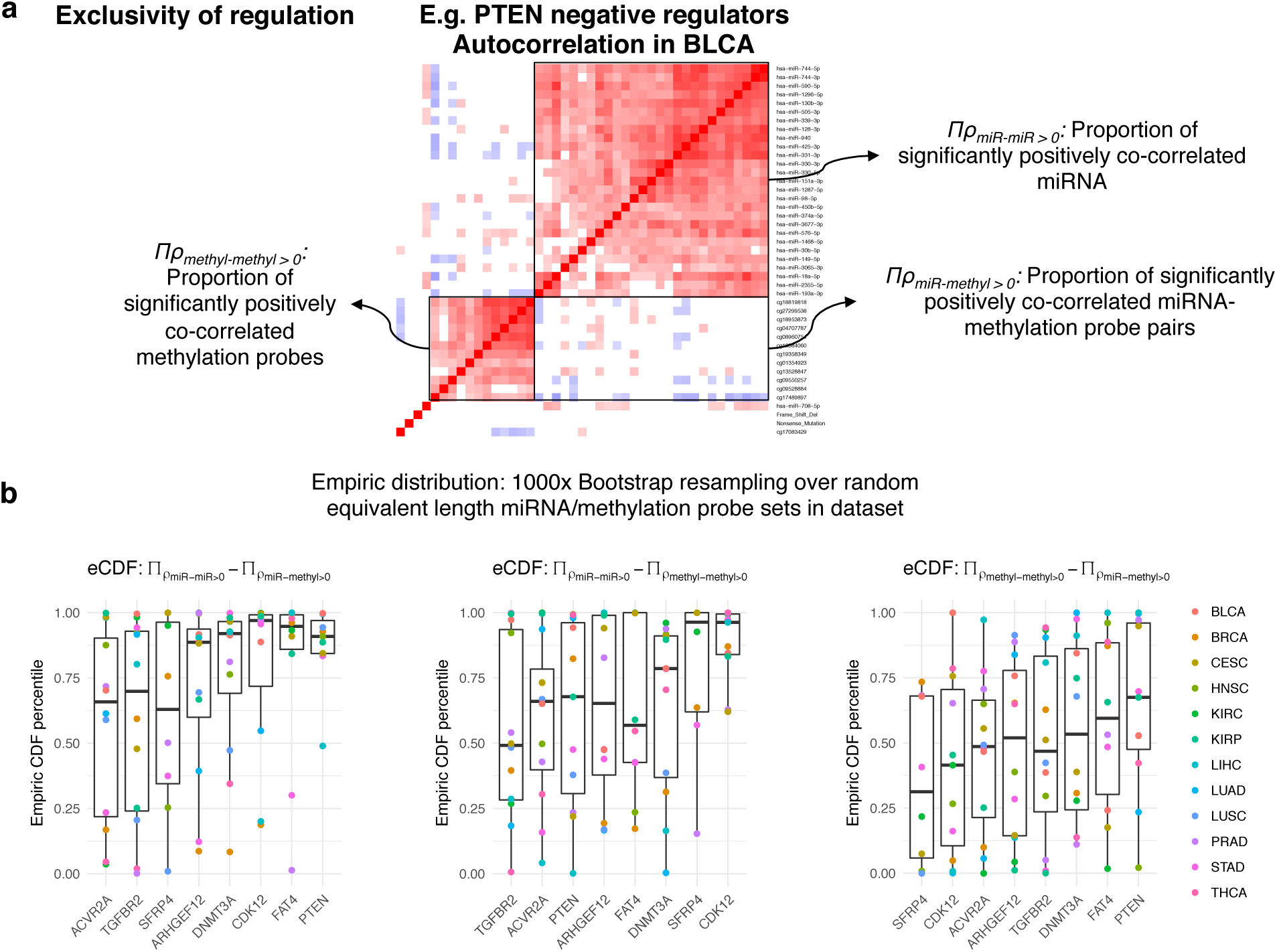
The approach used to determine the exclusivity of each mode of gene regulation on expression for the TSG considered. (**a**) Depiction of the autocorrelation heatmap for the expression of the various negative regulators of the tumour suppressor gene, and the variables considered and their meaning, as depicted. (**b**) Plots depicting the spread of the percentiles on the empiric CDF for the distributions for the pairwise differences of the variables identified in (**a**) through a bootstrapping-based analysis, as described in the Methods section.

### ARHGEF12, SFRP4, TGFBR2, and their cognate miRNAs, are consistently associated with breast cancer molecular subtype

Next, we sought to identify associations with tumour molecular subtypes, and as an initial analysis chose the molecular subtypes of breast cancer, owing to both the well-defined subtypes and the relatively large number of cases available for each subtype. An analysis of the eight identified tumour suppressor genes consistently negatively downregulated by miRNA across cancer types shows that in many cases, their mRNA levels are inversely associated with breast cancer molecular subtype. In particular, the basal subtype shows the lowest median expression of ARHGEF12, SFRP4, and TGFBR2, as compared to normal tissue, luminal A, B, Her2 amplified, or normal subtypes of breast cancer as shown in Supplementary Figure **??** in Section S11, and this association is retained when cases are restricted to wildtype expression of ARHGEF12, SFRP4, and TGFBR2. At the level of the associated miRNA identified as negative regulators of these TSG, we show that the median expression of these miRNA is also significantly associated with breast cancer molecular subtype, and inversely related to TSG mRNA expression by subtype. We have also shown that these associations are preserved when samples with non-silent mutations in the TSG are removed. For further validation, we also show reproducibility of these TSG and miRNA associations to breast cancer subtype in the independent Metabric dataset (*N* = 1293) [13].

## Discussion

In this work we have carried out a comprehensive and rigorous association analysis of human transcriptomic and genomic data to leverage an understanding of the role of miRNA in regulating complex phenotypes, through the lens of established gene expression signatures. Gene signatures represent transcriptomic association and we utilised them in two key ways, adding significant power to the analysis. Firstly, we use gene signatures to understand the relationship between non-coding RNA and phenotype; this exploits the phenotypic associations intrinsic to established gene signatures. Secondly, because miRNA can only repress mRNA that are present in sufficient quantity in a cell, when inferring function, it is vital to ‘group’ transcriptomic profiles by miRNA targeted gene expression. This allows for an understanding of the miRNA-mediated gene regulation important to the phenotype one wishes to uncover. Thus, this analysis represents a novel and powerful assessment of the complexity of miRNA regulation of phenotypes, particularly in the context of cancer.

Our work begins with ensuring applicability of the gene signatures, and then for each signature, we gain an understanding of the miRNA both significantly up - and down-regulated in association with the signature score. From this, we obtain the network shown in Figure 1, which describes for the first time in a detailed fashion, and across cancer types, the contribution of individual miRNA to the complex cancer phenotype. We also show reproducibility of this network in an independent dataset, by considering the overlap with the network reconstructed using the Metabric dataset and the same gene signatures. Moreover, repeating this analysis, grouping the miRNA significantly upregulated and down-regulated by miRNA family, illustrates that many miRNA families participate with members antagonistically across the hallmarks of cancer; including 4 of the top 5 most common miRNA families identified by our analysis (miR-25 family, miR-17 family, miR-15abc family, and let-7 family). This challenges the biological hypothesis of miRNA families acting in a generally coordinated fashion across multiple phenotypic states, and highlights the context dependent behaviour of individual miRNA themselves, regardless of grouping by family [20,28,29]. Further strengthening the argument for context-dependent actions of miRNA is the observation that we have made for the gene signature network reconstructed from 6 tissue types with samples of adjacent normal, non-tumour tissue. While a significant proportion (54%) of miRNA found to be associated with the gene signatures are the same as for the tumour tissues, the analysis of the targets of these miRNA reveals that they do not show enrichment for TSG in their targets, despite being concordant to the findings in tumour tissue, again highlighting the context dependency in miRNA-mediated gene regulation.

As might be expected, given the complexity of the action of non-coding RNA, we show in this work that for a given phenotype, single miRNA-target interactions do not account for the observed behaviour; rather it is subtle changes by a network of miRNAs, interacting with a set of targets in a coordinated manner, that serve to tune the transcriptome to achieve the complex phenotype. That is, because the targets of a given miRNA are predicted to be variable in their function, and are not all present in every sample at ‘repressable’ concentrations, the same miRNA can be associated with opposing phenotypic effects in different contexts, as reported by Denzler et al. in [14] for competing endogenous RNA. We show that the behaviour of miRNA is highly context dependent, and through the pan-cancer analysis, we have aimed to reduce the complexity of this context dependency by only selecting those interactions significantly occurring across cancer types. However, we caution that because miRNA are so context dependent, sample purity arises as an important issue in identifying pan-cancer miRNA signals. Further study into deconvolution methodologies enabling more accurate quantification of miRNA abundance from purely tumour samples will likely elucidate a clearer picture of miRNA-target interactions.

As miRNA are increasingly also thought of as potential therapeutic agents, because miRNA effects are highly context dependent and miRNA act in coordinated networks, if miRNA are to have effective therapeutic function, a single miRNA may be an ineffective strategy. Rather, we pose that a cocktail of miRNA will be necessary to sufficiently modify the tune of the symphony playing within the cancer cell, perhaps explaining poor therapeutic efficacy with current single miRNA-based therapeutics. For miRNA therapeutics to achieve function, we pose that these will likely have to be based on a number of miRNA, given to a highly selected group of patients with transcriptomes deemed to be responsive to this network perturbation, and that in patients without these profiles, such a cocktail would require modification in order to be effective. Further, by using more than a single miRNA as a therapeutic agent, the off-target effects that have significantly limited development in this field may be mitigated, by buffering for this with other miRNA in off-target tissues [1].

In this work we further the knowledge of which miRNA are involved in creating the phenotypes of cancer, across tissue types, to identify miRNA-TSG targets showing exclusive miRNA-mediated suppression. This suggests that a phenomenon similar to that of the previously described ‘BRCA-ness,’ wherein a miRNA, miR-182, has been shown to repress BRCA and confer sensitivity to PARP inhibitors in a subset of tumours [43], may be at work within many cases, and across multiple tumour suppressor genes. Additionally, recent work has shown how ‘epimutations’ may result in aberrantly methylated sites that can recapitulate the phenotype of a mutated tumour suppressor such as DNMT3A in leukaemia [27]. This raises the suggestion that there are tumour suppressor genes for which a mutation is not requisite for inactivation, but rather, inactivation is achieved through miRNA-mediated repression or methylation-mediated repression alone. For the TSG we have identified, we have also shown (see Online Materials), that the TSG mutations are occurring independently of MYC amplification status, which has been recently identified as an independent regulator of miRNAs. In addition, we show that such MYC amplification status is indeed associated with miRNA expression for the miRNA found to be negatively associated with each of the TSG in a majority of cases (Supplementary Figure **??**, Section S12). Further, we have shown that in particular tumours, for PTEN, CDK12, and FAT4, this miRNA or methylation-based suppression happens independently of other gene regulatory factors, such as mutations and copy number changes.

Lastly, we show how using generally validated, and specifically quality-controlled, gene signatures describing biologically conserved phenotypes can be used to collate large datasets to derive inference about miRNAs, a species whose signal has been traditionally hard to detect. The ability of this approach to capture tumour biology is highlighted through the identification of tumour suppressor genes showing miRNA-mediated regulation across tumour types, which we have shown have a very strong association to breast cancer molecular subtype. Specifically, this analysis points towards the role of decreased mRNA levels of ARHGEF12, SFRP4, and TGFBR2 in association with the poor-prognosis basal breast cancer subtype [2, 51]. Having identified potential negative regulators of these TSG, we show how these miRNA alone associate with breast cancer subtype, elevated in the basal subtype, capturing potentially novel biological association.

Finally, the presented methodology may be used in future work encompassing both more specific signatures, as well as larger, more expansive datasets to derive even greater confidence in particular associations. This approach will enable the functional annotation of a greater variety of miRNAs, illuminating their critical role in post-transcriptional gene regulation.

## Online Methods

### Gene signatures considered

We consider a wide variety of gene signatures, touching upon many of the hallmarks of cancer, as described in the original and updated work by Hana-han and Weinberg [24,25]. Signatures were selected through a review of MSigDB hallmarks signatures, as well as through a review of the literature, and those used are summarised in Table 3 [34]. We note that while many of these signatures were derived for a particular tumour type, we have applied them across many different tumour types, but before doing so, we have performed an evaluation step (*sigQC*) to ensure that each signature used is applicable to every dataset under consideration, in Supplementary section S1, Figures **??**- **??**.

### Datasets considered

In selecting datasets for this analysis, we initially aimed to select those comprising a comprehensive set of cancer types, with each type represented by a sufficient number of clinical samples, so as to reduce the effects of noise. Thus, we initially began with a consideration of all cancer types represented within the Cancer Genome Atlas datasets (TCGA), and limited based on origin of neoplasm and number of patients for whom miRNA-sequencing was carried out [55]. The RSEM normalised gene expression, mature miRNA normalised expression data, copy number, mutation, and methylation data were accessed from the Firebrowse database at http://www.firebrowse.org. In particular, we considered all cancer types which were epithelial or glandular with respect to histology, and with at least 200 samples with miRNA-sequencing data. These two filters limit the cancers considered to a total of 15 epithelial or glandular neoplasms, comprising a wide variety of cancer types, enabling the strong detection of fundamental biology. Furthermore, among these tumour types, there are 7,738 clinical samples, for which 7,316 have miRNA-sequencing data. The tumour types, along with their sample counts are presented in Table 1. Details of the number of samples included for each data type are presented in Table 2, and we note that for any analysis presented, any dataset present with fewer than 9 samples was excluded from analysis. This restriction excluded the analysis of COAD, OV, and UCEC datasets from the analysis of tumour suppressor genes, oncogenes, and exclusivity of regulation.

**Table 1.** **TCGA datasets considered and associated total clinical sample counts**.

| Dataset | Abbreviation | Clinical samples |
| --- | --- | --- |
| Breast invasive carcinoma | BRCA | 1098 |
| Ovarian serous cystadenocarcinoma | OV | 602 |
| Lung adenocarcinoma | LUAD | 585 |
| Uterine corpus endometrial carcinoma | UCEC | 560 |
| Kidney renal clear cell carcinoma | KIRC | 537 |
| Head and neck squamous cell carcinoma | HNSC | 528 |
| Lung squamous cell carcinoma | LUSC | 504 |
| Thyroid carcinoma | THCA | 503 |
| Prostate adenocarcinoma | PRAD | 499 |
| Colon adenocarcinoma | COAD | 460 |
| Stomach adenocarcinoma | STAD | 443 |
| Bladder urothelial carcinoma | BLCA | 412 |
| Liver hepatocellular carcinoma | LIHC | 377 |
| Kidney renal papillary cell carcinoma | KIRP | 323 |
| Cervical squamous cell carcinoma and endocervical adenocarcinoma | CESC | 307 |

**Table 2.** **Counts of common samples with miRNA, mRNA, mutation, methylation, and copy number data.**

| Dataset | mRNA samples | miRNA | mRNA and miRNA | mRNA, miRNA, mutation, methylation, and copy number |
| --- | --- | --- | --- | --- |
| BRCA | 782 | 755 | 499 | 324 |
| OV | 307 | 461 | 291 | 0 |
| LUAD | 517 | 452 | 449 | 181 |
| UCEC | 177 | 412 | 174 | 4 |
| KIRC | 534 | 255 | 255 | 121 |
| HNSC | 520 | 486 | 478 | 244 |
| LUSC | 501 | 342 | 342 | 51 |
| THCA | 501 | 502 | 500 | 396 |
| PRAD | 497 | 494 | 493 | 329 |
| COAD | 286 | 221 | 221 | 0 |
| STAD | 415 | 389 | 370 | 230 |
| BLCA | 408 | 409 | 405 | 128 |
| LIHC | 373 | 374 | 369 | 186 |
| KIRP | 291 | 292 | 291 | 148 |
| CESC | 304 | 307 | 304 | 190 |

**Table 3.** **Gene signatures considered and associated hallmarks of cancer.**

| Signature name | Reference | Number of genes | Associated hallmarks |
| --- | --- | --- | --- |
| Epithelial Mesenchymal Transition, MSigDB | MSigDB [34] | 200 | Activating invasion and metastasis |
| Invasiveness | Marsan et al., 2014 [38] | 16 | Activating invasion and metastasis |
| Oxidative Phosphorylation, MSigDB | MSigDB [34] | 200 | Deregulating cellular energetics |
| Reactive Oxygen Species Pathway, MSigDB | MSigDB [34] | 49 | Deregulating cellular energetics |
| G2M Checkpoint, MSigDB | MSigDB [34] | 200 | Enabling replicative immortality |
| PI3K-AKT-MTOR Signaling, MSigDB | MSigDB [34] | 105 | Evading growth suppressors |
| Xenobiotic Metabolism, MSigDB | MSigDB [34] | 200 | Evading growth suppressors |
| DNA Repair, MSigDB | MSigDB [34] | 150 | Genome instability and mutation, Enabling replicative immortality |
| p53 Pathway, MSigDB | MSigDB [34] | 200 | Genome instability and mutation, Enabling replicative immortality |
| Hypoxia | Buffa et al., 2010 [5] | 51 | Inducing angiogenesis |
| Angiogenesis, MSigDB | MSigDB [34] | 36 | Inducing angiogenesis |
| Hypoxia, MSigDB | MSigDB [34] | 200 | Inducing angiogenesis |
| Angiogenesis, upregulated | Desmedt et al., 2008 [15] | 5 | Inducing angiogenesis |
| Angiogenesis | Masiero et al., 2013 [39] | 43 | Inducing angiogenesis |
| Proliferation, upregulated | Desmedt et al., 2008 [15] | 140 | Sustaining proliferative signaling |
| KRAS Signaling, Up, MSigDB | MSigDB [34] | 200 | Sustaining proliferative signaling |
| Inflammatory Response, MSigDB | MSigDB [34] | 200 | Tumour-promoting inflammation, Avoiding immune destruction |
| IL2-STAT5 Signaling, MSigDB | MSigDB [34] | 200 | Tumour-promoting inflammation, Avoiding immune destruction |
| IL6-JAK-STAT3 Signaling, MSigDB | MSigDB [34] | 87 | Tumour-promoting inflammation, Avoiding immune destruction |
| TGFβ Signaling, MSigDB | MSigDB [34] | 54 | Tumour-promoting inflammation, Avoiding immune destruction |
| TNFα Signaling via NF-κB, MSigDB | MSigDB [34] | 200 | Tumour-promoting inflammation, Avoiding immune destruction |
| Immune Invasion, upregulated | Desmedt et al., 2008 [15] | 92 | Tumour-promoting inflammation, Avoiding immune destruction |

### miRNA family database

miRNA ranked across different cancer types were further grouped together by microRNA family, as defined by the targetscan database, implemented in R as the targetscan.Hs.eg.db package [12,33].

### Statistical methodology

#### Transcriptomic data

Data were taken from the GDAC Firebrowse TCGA portal provided by the Broad Institute. miRNA datasets used were log2 normalised mature miRNA counts for all cancer types. mRNA datasets used were normalised RSEM genes taken from data through the Illumina HiSeq RNAseq v2 platform. These expression data were then transformed by the transformation log_2_(*x* + 1), for *x* as the original expression value, and this was used in all further computation for all cancer types and signatures. Where not otherwise specified, signature scores are taken as the median of log2-transformed expression of all signature genes for each sample. Metabric datasets for normalised miRNA and mRNA expression were taken from the European Genome-Phenome Archive (EGA) under study accession numbers EGAD00010000434 and EGAD00010000438.

#### Penalised linear regression

The aim of the penalised linear regression methodology was to determine those miRNA which most strongly predict (positively or negatively), the gene expression summary score for each signature. With consideration of this, the linear regression was designed such that the model utilised the expression levels of each individual miRNA as a covariate, in order to predict the signature score, taken as the median of the log-transformed expression levels of the signature genes. We note that in order to facilitate direct comparability between distinct signatures and caner types, we first normalised both the scores and miRNA expression levels to a mean of zero and unit variance. This transformation ensures that the coefficients and their relative values are comparable between cancer types and signatures.

We used multivariate penalised linear regression, with 10-fold cross validation, as previously described [6] to infer significant relationships between miRNA and gene signatures without overfitting our model. Specifically, first a univariate model filter was applied to the data to select miRNA used for penalised multivariate linear regression. Then, the penalised multivariate linear model with the least predictive error (as assessed on the validating folds) was selected, and coefficients for these miRNA were used for further analysis. All model-fitting, including the initial filtering, was done with 10-fold cross-validation, and was carried out using the *penalized* package in R [21, 22]. The initial univariate filter was applied to remove miRNA showing little predictive power from the multivariate linear model, and only those miRNA with *p* < 0.2 significance in the univariate linear model predicting signature score were considered. This permissive p-value was used to assure that the multivariate linear model did not contain artificially stringent associations, as the penalization procedure also functions as a stringency filter, reducing the false discovery rate. The multi-variate linear regression was carried out as a penalised L1/L2 regression to reduce complicating effects of co-correlated miRNAs as predictors of the signature scores. To tune the parameters for the combined L1/L2 regression, a range of values (0, 0.01, 0.1, 1, 10, 100), was tested for the L2 parameter, while in each case the L1 parameter was optimised. Following computation of all models, the model with the greatest log-likelihood was chosen.

#### Rank product analysis

Once coefficients were obtained for the linear model via the penalised regression approach described earlier, these were collated into matrices with columns defined by cancer type, for each of the gene signatures considered. These coefficients were then fractionally-ranked both from most negative to most positive, and most positive to most negative in value. The rank product statistic, as described by Breitling et al., in 2004, for these fractional ranks was then considered, and the coefficients were ranked in terms of their significance of rank product test statistic, as implemented by the RankProd R package [4,9]. This was used to give high-confidence rankings of miRNA associated both positively and negatively with the various signatures considered.

### Validation of miRNA-signature interactions

In order to ensure reproducibility of the approach used to identify gene signature-associated miRNA, we repeated the linear modeling procedure across the independent Metabric matched miRNA and mRNA microarray dataset of 1293 samples [13]. We mapped each gene signature to corresponding Ensembl IDs, and repeated the combined univariate-multivariate linear modeling approach over all miRNA probes. The miRNA probes identified as positive and negative coefficients were then identified, and mapped to their corresponding mature miRNA ID. The statistical significance of this overlap is shown in Supplementary Figure **??**, and was calculated using the Fisher exact test. Nearly all signatures show strong statistical significance, and in the majority of cases not reaching statistical significance, signature applicability to the Metabric dataset may present an issue, as signatures contained a high proportion of genes with low variance, which presents an issue for signature applicability, particularly for microarray-based datasets.

### Target analysis

Targets were aggregated for each miRNA using the miRNAtap database in R, as implemented through the Bioconductor targetscan.Hs.eg.db package [46]. The default settings of using all 5 possible target databases: DIANA [37], Miranda [17], PicTar [32], TargetScan [19], and miRDB [56], with a minimum source number of 2 were used, and the union of all targets found was taken as the set of targets for a given miRNA.

For each of these target-miRNA pairs, the Spearman correlation coefficient was calculated across every cancer type for miRNA versus target mRNA expression, partial to mutation status of the mRNA, and if this value reached statistical significance of *p* < 0.05, it was recorded, and otherwise was omitted and recorded as NA. Note that mutational status was reported as a binary variable with a value of 1 for any non-silent, non-intronic mutation, and 0 otherwise. The target-miRNA pairs with at least 5 non-zero entries across cancer types were kept for further analysis, and subsequently were analysed using the rank product statistic, to identify those pairs with consistently negative correlations, across cancer types, with respect to all other hallmarks-miRNA pairs. Partial correlations were done in R using the ppcor package [30].

Furthermore, in the global analysis of all TSG-miRNA pairs, we considered every TSG-miRNA predicted target pair, and again considered the Spearman correlation partial to mutation status, omitting the value as NA if significance *p* < 0.05. The rank product statistic was again considered on those pairs with at least 5 non-zero values across cancer types, thereby identifying those TSG-miRNA pairs consistently negatively correlated across cancer types, significantly with respect to all other TSG. Lists of known oncogenes and tumour suppressor genes were taken from the COSMIC database [18]. Because MYC amplification is a possible confounder to the miRNA identified as associated with TSG across cancer types, we checked to ensure that mutation of the 8 TSG identified, across cancer types, does not co-occur significantly with MYC amplification. Of the 96 TSG-cancer type pairs (8 TSG over 12 cancer types), none showed significance in the over-enrichment by a one-sided Fisher exact test for MYC amplification and TSG mutation after correcting for multiple testing.

### Bootstrap resampling to determine significance of TSG enrichment among repressed targets

To determine the significance of the number of the TSG repressed among the repressed targets of the miRNA identified as signature-associated, we resampled from all miRNA that could possibly selected as signature associated (i.e. those with at least 80% non-zero expression across samples in at least one tumour type), and created 1000 resampled lists of random miRNA of the same length as the number of signature-associated miRNA. Using these lists and the methodology above, miRNA targets were identified, and those miRNA-target correlations (partial to mutation status) consistently negatively ranking compared to all others across tumour types were recorded for each list. Among these repressed targets, we identified the number of TSG overlapping with the COSMIC TSG list, and used this to define the empirical distribution of the number of TSG overlapping with the miRNA targets. Then from this distribution, to determine the significance for the 21 TSG overlapping the repressed TSG targets of the signature-associated miRNA, we determined the empirical CDF percentile for the value 21, reported as 0.983, yielding *p* = 0.017 from this analysis. To ensure that 1000x bootstrap resampling was sufficient, we used the QQ plot for the empirical distribution to ensure close adherence to normality for this distribution.

### Analysis of TSG regulation

In analysing the regulation of the TSG identified as related to the hallmarks of cancer and potentially amenable to miRNA regulation, we first limited the samples under consideration to those where copy number data, gene expression data, miRNA expression, mutation data, and methylation data were all present. Mutation data was again taken as a binary variable, but as opposed to the partial correlation analysis, mutations were stratified into their reported types (e.g. missense mutations are all grouped together, etc.). That is, the missense mutation variable would only contain a value of 1 if the sample had a missense mutation in the gene of interest, and 0 otherwise. All variables considered in the linear regression were standardised to a mean of 0, and a standard deviation of 1.

L1/2 penalty-based penalised linear regression was then performed, in the same manner as above, for the linear model described in Figure 3a. Subsequently, coefficients were aggregated across the various cancer types and after the rank product test was applied, those predictors showing statistically consistent positive or negative coefficients were identified. Following this, the autocorrelation of each of these predictor variables was considered, for each of the TSG in each cancer type, as depicted by the heatmap in Figure 4a.

### Analysis of the exclusivity of gene regulation

To determine the exclusivity of gene regulation, we calculated the empiric distributions of the variables Π_*Pk*_ as defined graphically in Figure 4. These represent the proportion of miRNA-miRNA or miRNA methylation or methylation probe-methylation probe pairs that show significant positive Spearman co-correlation (*p* < 0.05). For the bootstrapping analysis, we resampled the datasets, choosing miRNA and methylation probes in the same number as the heatmap in question, and then considered the distributions of the pairwise differences in the variables Π_*Pk*_. From these distributions for the pair-wise differences, we were able to infer the percentile on the empirically constructed CDF that the true case represented, the results of which are depicted in Figure 4b, showing, for each gene and cancer type, the percentile on the pairwise difference empiric distribution for the observed heatmap.

The calculations for the analysis of TSG regulation and analysis for the exclusivity of gene regulation were repeated for an idependent dataset comprising matched mRNA, miRNA, CNV, mutation, and methylation data for 93 patients with ovarian cancer, from the OV-AU project from the ICGC data portal [58]. Results of this analysis are highlighted in Supplementary Section S9, Figure**??**.

## Acknowledgements

The authors are grateful for the financial support provided by Cancer Research UK to FMB, ALH and AD, and the support provided by the Clarendon Fund to AD. ALH is grateful for the financial support provided by the Breast Cancer Research Foundation. The authors thank Dr. Venkata Manem for helpful discussions. All authors are eternally grateful to the patients and their families who provided tissue to the Cancer Genome Atlas and Metabric projects.

## Author Contributions

FMB conceived the idea, designed and supervised the study. AD and FMB devised the analyses. AD wrote code and performed analyses. JGS, ALH and FMB supervised the implementation. AD wrote the manuscript with contribution from all other authors.

